# Learning and embodied decisions in active inference

**DOI:** 10.1101/2024.08.18.608439

**Authors:** Matteo Priorelli, Ivilin Peev Stoianov, Giovanni Pezzulo

## Abstract

Biological organisms constantly face the necessity to act timely in dynamic environments and balance choice accuracy against the risk of missing valid opportunities. As formalized by embodied decision models, this might require brain architectures wherein decision-making and motor control interact reciprocally, in stark contrast to traditional models that view them as serial processes. Previous studies have assessed that embodied decision dynamics emerge naturally under active inference – a computational paradigm that considers action and perception as subject to the same imperative of free energy minimization. In particular, agents can infer their targets by using their own movements (and not only external sensations) as evidence, i.e., via *self-evidencing*. Such models have shown that under appropriate conditions, action-generated feedback can stabilize and improve decision processes. However, how adaptation of internal models to environmental contingencies influences embodied decisions is yet to be addressed. To shed light on this challenge, in this study we systematically investigate the learning dynamics of an embodied model of decision-making during a *two-alternative forced choice* task, using a hybrid (discrete and continuous) active inference framework. Our results show that active inference agents can adapt to embodied contexts by learning various statistical regularities of the task – namely, prior preferences for the correct target, cue validity, and response strategies that prioritize faster or slower (but more accurate) decisions. Crucially, these results illustrate the efficacy of learning discrete preferences and strategies using sensorimotor feedback from continuous dynamics.

## 1 Introduction

The study of value-based decision-making in psychology and neuroscience has often focused on static settings, in which participants are asked to choose between a fixed number of choice alternatives (usually two) on the basis of known attributes, such as probabilities, utility, temporal delay, or their combinations. These situations are generally characterized in terms of a family of serial (*decide-then-act*) models, in which a decision is firstly made upon the accumulation of sensory evidence to a threshold and then reported by acting, e.g., by pressing a response button. Serial models such as the drift-diffusion model, the leaky, competing accumulator model, or the race model have been highly successful in explaining a wide variety of human behavioral and neural data, during value-based and perceptual decisions [36,41].

Living organisms, however, often face *embodied decisions* that differ significantly from the value-based decisions widely studied in the laboratory [7,20,44]. Consider a lion chasing gazelles in the savanna or a driver surpassing other drivers while avoiding collisions. The embodied character of the decisions is mostly evident from the fact that agents make choices not just about potential outcomes but also about potential action plans to achieve them – or between competing affordances for movement, i.e., an affordance competition process [8,5,27]. Environments can also change rapidly and require updating of action plans on the fly. Thus, agents must implement these plans promptly to avoid losing valued opportunities. Differently from static decisions studied in the laboratory, embodied decisions can regard an open-ended number of choice alternatives and features that are not necessarily defined (or known) a priori and that might be continuous and change over time.

Various empirical studies addressed embodied decisions, both in minimally embodied setups in which classical decision tasks are augmented to require simple action dynamics, and in more sophisticated setups that mimic more closely the competition between movement affordances faced by animals in their lives [4,14,38,42]. These studies reveal that the serial (*decide-then-act*) view is insufficient to fully account for embodied decisions, for two main reasons. First, action and decision dynamics can unfold in parallel, in such a way that movement dynamics provide a rich readout of the ongoing decision [13,39]. Second, and perhaps more importantly, action dynamics can influence choice, e.g., in terms of motor costs associated with the alternatives [24,2,10,22].

Some of these findings are well accounted for by a novel class of embodied decision models that go beyond serial assumptions. For example, the *affordance competition* model assumes that the brain can specify, evaluate, prepare and sometimes even execute potential action plans in parallel [8,5,27]; in turn, this might require distributing the burden of decision-making across several circuits and networks, therefore implementing choice as a distributed consensus rather than as a centralized process [6]. Other embodied decision models also incorporate feedback from action to decision-making, simultaneously optimizing decision and action processes [4,23]. A recent computational study [35] showed that embodied decision dynamics emerge naturally under active inference, a computational paradigm that considers action and perception as subject to the same imperative of free energy minimization [26]. Key to this model is the reciprocal loop between *motor planning* – during which beliefs about the target to be reached contribute to selecting an appropriate motor plan – and *motor inference* – during which target-directed plans and movements are used as evidence to update beliefs about the target to be reached, in parallel to sensory evidence. Under appropriate conditions, this continuous interaction can stabilize and improve decisions.

However, that study was based on a fixed generative model of the task and left unaddressed the way agents can learn and update their models. Humans and other animals show robust learning of the statistical regularities of cognitive tasks, adapting their strategies and response dynamics over time. For example, during the Flanker [21] and Posner tasks [29], it is possible to learn the probability of the correct response or the probability that some cues predict the correct response (called *cue validity*) within a certain experimental block. In these tasks and many others, participants learn expectations about targets, cues or other elements, which influence their responses and movements in subsequent trials [43,19].

For these reasons, in this study we systematically investigate the learning dynamics of an embodied model of decision-making during a two-alternative forced choice task, by relying on a hybrid (discrete and continuous) active inference framework [35]. Our results show that the active inference agent can not only learn various statistical regularities of the task – namely, prior preferences for the correct target or cue validity – but also those characteristics peculiar to embodied models of cognition, i.e., the response strategies that prioritize faster or slower (but more accurate) decisions.

## 2 Methods

### 2.1 The two-alternative forced choice (2AFC) decision task

To study learning in embodied decisions, we designed a two-alternative forced choice (2AFC) decision task with time-varying information, i.e., wherein evidence for one choice or the other, expressed in terms of sequentially provided cues, changes throughout each trial (Figure 1a). The agent’s body is a 3-DoF arm starting from a position at an equal distance from the two target buttons (red and green circles), and has to reach the target that will contain more cues (the smaller grey dots) in it. During the task, one cue after the other appears either in the left or the right circle (15 in total). The agent can move at any moment, until a certain deadline. By varying systematically the probability distributions from which the cues are sampled, we create two types of conditions: easy conditions in which cues appear in the correct target with an initial probability of 80 %, which then gradually increases to 100 % after 8 cues (*congruent* trials); and difficult conditions in which cues appear in the correct target with an initial probability of 20 % which then increases to 100 % (*incongruent* trials). Hence, during incongruent trials the cues will initially appear in the wrong target. The correct target (i.e., the circle that will contain the most cues) is sampled randomly at each trial.

**Fig. 1:**
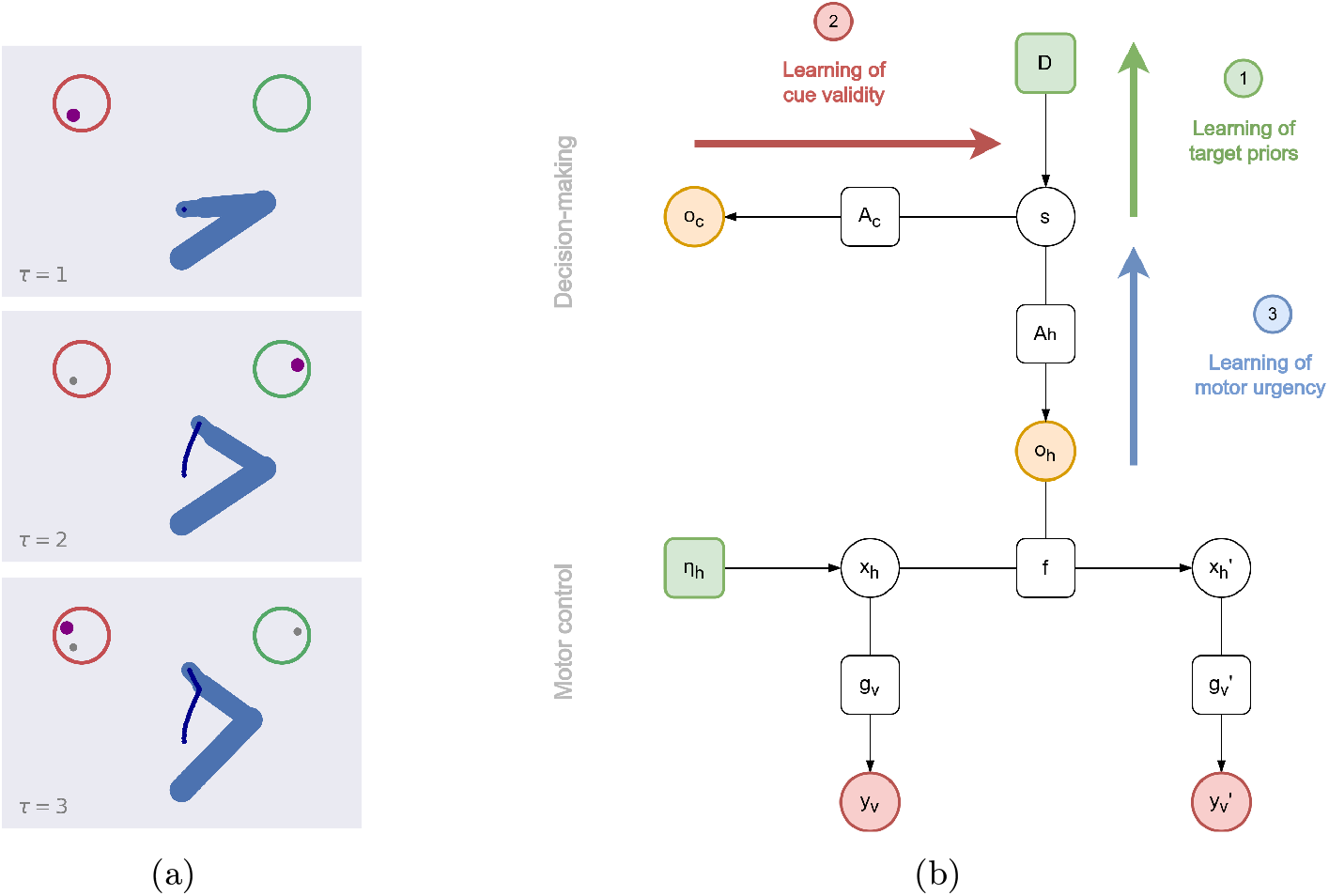
Embodied decision setup and active inference model. (a) Experimental setup, during three consecutive discrete time steps *τ*. The agent controls a 3-DoF arm, which starts at a home position (blue dot) at an equal distance from the two targets (red and green circles). The current cue is displayed with a big purple dot, while the old cues are shown with smaller grey dots. For each trial, the agent has to reach the target it believes will contain more cues. (b) Factor graph of a hybrid active inference model for embodied decisions. Variables and factors are indicated by circles and squares, respectively. We highlighted three pathways related to the learning of the correct target (green arrow), cue validity (red arrow), and response strategy (blue arrow).

The 2AFC task used here addresses the most important features of classical decision tasks (e.g., it needs making a choice based on sensory evidence, which can have different sequential statistics) but in a minimally embodied setting, as it requires a reaching movement to respond. The 2AFC task is similar to the Tokens task designed in [9], except for the fact that only a single cue at a time is visible by the agent, and that action starts from a point below the targets, not in between. Furthermore, the 2AFC task presents some analogies with the classical *Eriksen flanker task* used to analyze attention mechanisms and assess the ability to suppress inappropriate responses [12]. In the flanker task, participants observe a visual target along with congruent or incongruent cues.

The change in the probability distributions from which the cues are sampled in our task is conceptually similar to the progressive shift of attention toward the correct target typically observed in the flanker task.

### 2.2 Hybrid active inference model for embodied decisions

To address the 2AFC task, we implemented a hybrid active inference model – shown in Figure 1b – composed of two parts: a discrete model that accumulates evidence over the cues and infers the correct target, and a continuous model that deals with the actual motor execution to reach it (see also [35]).

The discrete hidden states ***s***, encoding the probability that each target is the correct choice for the current trial, are sampled from a categorical distribution, i.e., ***s*** = *Cat*(***D***) = [*s*_*t*1_ *s*_*t*2_], where ***D*** are the parameters of a Dirichlet distribution and define the agent’s prior beliefs. These are iteratively inferred from discrete cues ***o***_*c*_ by inverting the cue likelihood matrix ***A***_*c*_. This matrix takes into account some uncertainty *α*_*c*_ over the cues with a similar role to the *drift rate* in driftdiffusion models:

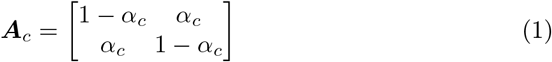

To compute a motor plan, the hidden states ***s*** also generate a particular combination of hand dynamics ***o***_*h*_ through the likelihood matrix ***A***_*h*_. Specifically, ***o***_*h*_ encodes the probability of reaching the left target, reaching the right target, or staying, i.e., ***o***_*h*_ = [*o*_*h*,*t*1_, *o*_*h*,*t*2_, *o*_*h*,*s*_]. A parameter *α*_*h*_ controls the weight of the last dynamics:

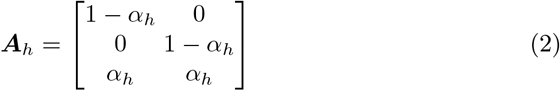

Note that the lower *α*_*h*_, the less certain the agent has to be about the correct target to start moving. At each discrete step *τ*, a particular cue ***o***_*c*_ and a particular hand dynamics ***o***_*h*_ are observed and compared with the corresponding predictions. Then, the inference of the discrete hidden states ***s*** follows the equation:

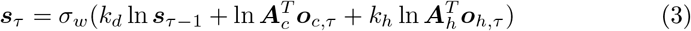

where *k*_*d*_ and *k*_*h*_ are scale parameters, and *σ* is a weighted softmax function, whose precision ensures fast transitions between discrete states, promoting less uncertain decisions. In all simulations, we set *k*_*d*_ = 1.0. In Equation 3, we note a prior coming from the previous step (equal to ***D*** at the beginning of the trial), a sensory likelihood for evidence accumulation, and a likelihood linked to the hand dynamics. The latter behaves as a sensory signal for the discrete model (similar to ***o***_*c*_) and causes the agent to infer the correct target through its own movements. This *self-evidencing* mechanism of motor inference stabilizes the decision taken [1], a behavior observed in many biological scenarios [23]. See [35] for more details.

The discrete set of hand dynamics ***o***_*h*_ is inferred via Bayesian model comparison, i.e., by comparing a prior surprise encoding the dynamics generated by the discrete model based on its guess, and a log evidence encoding the most likely hand dynamics corresponding to the current motor trajectory:

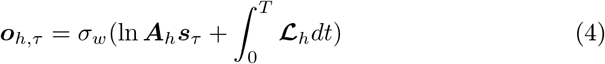

For each discrete step *τ*, the log evidence is accumulated over a continuous period *T*. The derivation of Equation 4 from the free energy associated with each discrete observation can be found in [18,25]. Generating predictions about hand dynamics (as opposed to positions as in conventional hybrid models) allows the agent to interact with dynamic elements of the environment. As before, *σ* _*w*_ is a weighted softmax whose precision controls how fast the transition between different dynamics occurs – e.g., high and low precisions are respectively related to abrupt and gradual movement onsets. Each hand dynamics *o*_*h*,*m*_ is linked to a continuous dynamics function, or potential motor plan ***f***_*m*_ in extrinsic (e.g., Cartesian) coordinates:

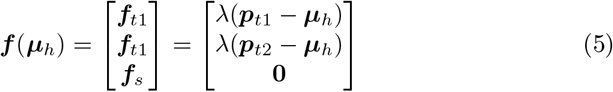

where ***μ*** _*h*_ is the belief over the hand position ***x***_*h*_, λ is an attractor gain, while ***p***_*t*1_ and ***p***_*t*2_ are the positions of the two targets – assumed to be known and fixed. The log evidence *ℒ*_*h*,*m*_ scores how much the *m*th potential dynamics is close to the belief 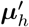 over the real dynamics perceived by the agent:

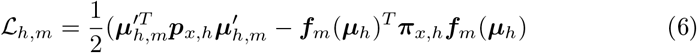

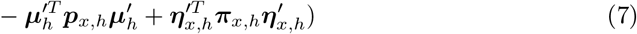

where ***f***_*m*_(***µ***_*h*_) is the *m*th dynamics function with precision ***π***_*x*,*h*_, the posterior 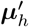 encodes the estimated velocity with precision ***p***_*x*,*h*_, and 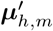 is the posterior of the *m*th dynamics. See [17,16] for more details about Bayesian model reduction, and [31,34] regarding hybrid models in dynamic contexts.

The motor plan to be realized is instead computed via Bayesian model average, i.e., by weighting each dynamics function with the respective discrete probability, i.e.,

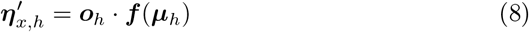

Hence, 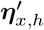 represents an average trajectory that accounts for the probability of each potential dynamics based on some discrete goal. This desired velocity enters the update of the continuous hidden states as a dynamics prediction error 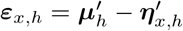. The continuous hidden states encode the hand position and velocity in extrinsic coordinates, and are updated – using the generalized beliefs 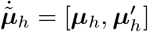 – through the following rule:

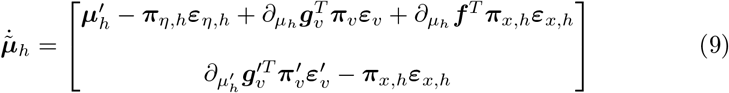

Here, we note two likelihood terms coming from visual observations ***y***_*v*_ and 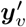 for both orders which invert the respective generative models and keep the belief close to the actual hand trajectory; the dynamics prediction error ***ε***_*x*,*h*_, affecting both orders either as a forward or backward message; and a prior prediction error ***ε***_*η*,*h*_ that biases the belief over the hand position. The latter comes from another continuous model encoding proprioceptive trajectories (e.g., expressed in joint angles) and performing forward kinematics. As a result, the backward message automatically performs inverse kinematics, eventually driving action. See [32,30] for more details about kinematic inference.

## 3 Results

Here, we describe three simulations addressing different aspects of the learning dynamics of our embodied active inference agent during the 2AFC task. The three learning processes are illustrated in Figure 1b; these comprise learning the priors over the correct target (green arrow), cue validity (red arrow), and a response strategy (blue arrow). In the following, we separately analyze each of them, keeping the rest of the model parameters fixed.

### 3.1 Simulation 1: Learning priors over the correct target

Many cognitive tasks involve learning statistical regularities; in the 2AFC task, the most common is perhaps the probability (across trials) of the correct target. Here, we simulate this learning by relying on the Dirichlet priors (or simply, counts) ***d*** over the discrete hidden states ***s***, i.e., *p*(***D***) = *Dir*(***d***) [37,11,15,40]. In active inference, learning implies that after every trial, the counts associated with each target are updated as follows:

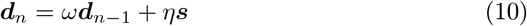

where *n* is the trial number, *ω* is a forgetting factor of older trials (usually initialized with a value reflecting high confidence over the prior belief), and *η* is the learning rate of new trials. Then, the counts are normalized to compute the priors ***D*** of the correct target for the successive trial. To assess the agent’s ability to learn the target distribution, we simulate 50 *incongruent* trials, with two phases. In the initial learning phase (first 10 trials), cues appear in the left target with a starting probability of 20%, which then gradually increases to 100%. In the second reversal learning phase (subsequent 40 trials), the condition is reversed so that the correct target is the green circle. The counts ***d*** are initialized to 0.5, while the forgetting and learning rates are set to *ω* = 0.99, *η* = 0.2.

The results of this simulation are shown in Figure 2b: during the early trials of the first phase (dark blue trajectories in Figure 2a), the agent moves toward the wrong direction and then changes mind. However, in later trials (dark red trajectories) it begins to move early toward the correct target, ignoring the accumulation of wrong cues. In parallel, movement onset decreases (Figure 2b). This result shows how a strong prior can overcome conflicting evidence. In the second phase, after the reversal at trial 10, the discrete prior for the left target slowly decreases, as the Dirichlet counts for the right target increase. In early trials, movement curvature increases and movement onset is slower, as the agent is uncertain about the distribution of the cues. In late trials, movement curvature decreases and movement onset fastens, as the agent learns the novel contingencies. This result shows that an embodied model can flexibly adapt to novel contingencies, solving reversal learning tasks.

**Fig. 2:**
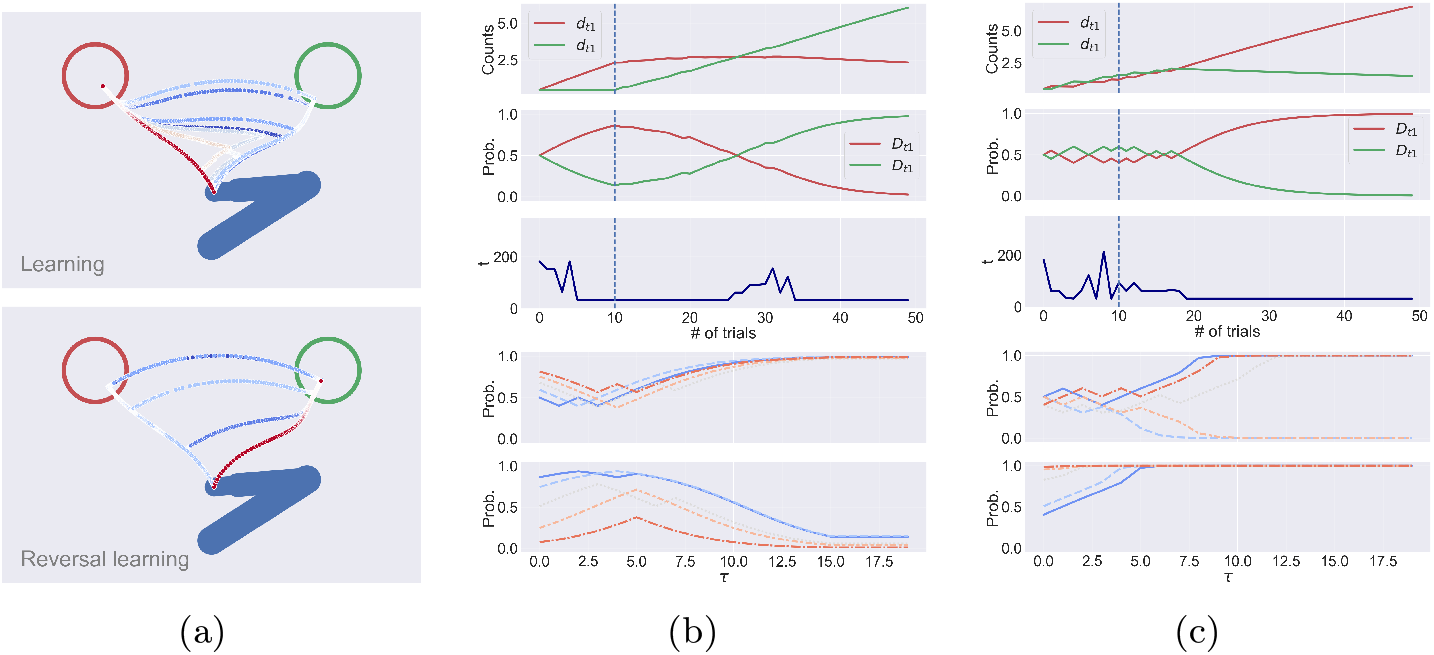
Results of Simulation 1. Statistical learning of the prior for the correct choice over 50 incongruent trials, composed of two phases in which the correct choice are the left and right targets, respectively. (a) Hand trajectories in equally spaced trials, during the first learning phase (top) and the second, reversal learning phase (bottom). Dark blue trajectories represent early trials, while dark red trajectories are late trials. Here, *k*_*h*_ = 0, *α*_*c*_ = 0.4, and *α*_*h*_ = 0.4. (b) The five panels show Dirichlet counts ***d***; discrete priors ***D***; time step of movement onset across trials; discrete hidden state *s*_1_ (which is associated with the left target) for learning; and reversal learning, in 5 equally spaced trials over discrete time *τ* on a blue-to-red color scale. The vertical dashed lines indicate the time step when reversal occurs. (c) Every parameter is the same as the previous simulation, but in this case *k*_*h*_ = 0.05.

Finally, Figure 2c shows statistical learning of the target priors in conditions identical to the previous simulations, but in this case the agent also exploits its motor responses to infer the correct choice, i.e., with *k*_*h*_ = 0.05 – resulting in a completely different behavior. As analyzed in [35], inferring the choice through one’s own trajectories reinforces the decision taken and stabilizes the action, resulting in fewer changes of mind and more confident behavior. This generally optimizes the speed-accuracy tradeoff and decreases the risk of losing valid opportunities in a dynamic environment. However, as we show in Figure 2c, the early presentation of incongruent cues during motor inference can lead to the formation of wrong habits. In fact, the cues initially appearing in the green circle reinforce the related decision, from which a habit of reaching the wrong target gradually emerges over several trials. This result might provide a hint regarding the emergence of pathological conditions in goal-directed behavior.

### 3.2 Simulation 2: Learning cue validity

In active inference, not only can statistical regularities over the hidden states be learned, but also the mapping from hidden states to sensory outcomes, i.e., the likelihood matrix ***A***. This form of learning is crucial in dynamic environments, where sensory uncertainty might change depending on the context. For example, during the Posner task [29], it is common to vary the validity of the cues (e.g., the probability that a cue predicts the correct response) across blocks. In more mundane situations, the repeated observation of high noise on a sensory modality (e.g., turning off the lights in a room) implies that the agent has to decrease its confidence about that modality and rely more on other clues (e.g. tactile or auditory sensations), and this should appear as increased uncertainty in the related likelihood matrices. This increase (or decrease) of confidence generally occurs when the agent’s predictions are repeatedly met (or violated).

The adaptation of the likelihood matrix ***A*** breaks down to keeping count of coincidences between states and outcomes. More formally, if we express ***A*** by the following Dirichlet distribution:

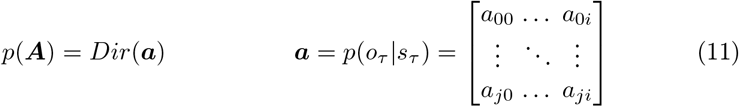

we can update the counts ***a*** similar to the learning of the prior ***D***:

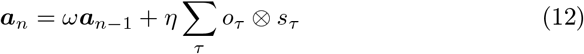

where *ω* and *η* are forgetting and learning rates, and ⊗ denotes the outer product (see [37] for more details). As evident, the only difference with Equation 10 is that learning is not based on the probability of the hidden states at the end of each trial, but on every occurrence of state-outcome pairs within trials. This kind of learning follows the Hebbian rule, according to which “neurons that fire together, wire together”. In fact, active inference assumes that the concentration parameters can be associated with the strength of synaptic connections [15].

In our task, variations in uncertainty over the cue mapping become evident when the difficulty of the trials abruptly changes. To show this, we ran 30 congruent trials (equivalent to a block of the Posner task in which the cue has high validity) followed by 30 incongruent trials (equivalent to a block of the Posner task in which the cue has low validity), and analyzed the agent’s behavior during learning of the likelihood matrix ***A***_*c*_. The counts ***a***_*c*_ are initialized with low values to provide a weak prior on evidence accumulation, *α*_*c*_ = 0.44:

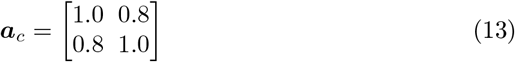

The forgetting and learning rates are respectively set to 0.98 and 0.01, while the parameter *α*_*h*_ controlling the strength of the stay dynamics is kept fixed to 0.4. Note that although we let the agent adapt to a generic form of the likelihood matrix, by sampling both targets as correct choices the initial parameterized form defined in Equation 1 is maintained (but in general this is not needed). As evident from Figure 3a, during congruent trials the parameter *α*_*c*_ (values on the antidiagonal) rapidly decreases as the agent’s prediction of the correct target matches almost every cue – a direct consequence of the accumulation of counts shown in the first column of Figure 3b. Instead, repeated exposure of more difficult (incongruent) trials leads to a slow decrease of *α*_*c*_, since the variability of the cues is much higher during the first half of the trials. This change of uncertainty reflects the rate of accumulation and hand velocity, as represented in the second and third columns of Figure 3b. In particular, the second column shows that during early congruent trials (displayed in blue), the discrete state ***s***_*t*1_ associated with the left target updates slowly, never reaching complete confidence; however, during late congruent trials (displayed in red in the top panel), the update of ***s***_*t*1_ is much more rapid, reflecting the increased confidence over cue validity. A specular behavior can be seen during incongruent trials (bottom panel), during which the rapid target estimation reflects the confidence learned in the previous congruent trials, which however slowly returns to the initial rate of accumulation as repeated difficult trials are observed. Regarding hand velocity, the third column of Figure 3b shows that the agent’s responses (here represented by the belief 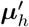) are gradually anticipated and increase in magnitude during congruent trials.

**Fig. 3:**
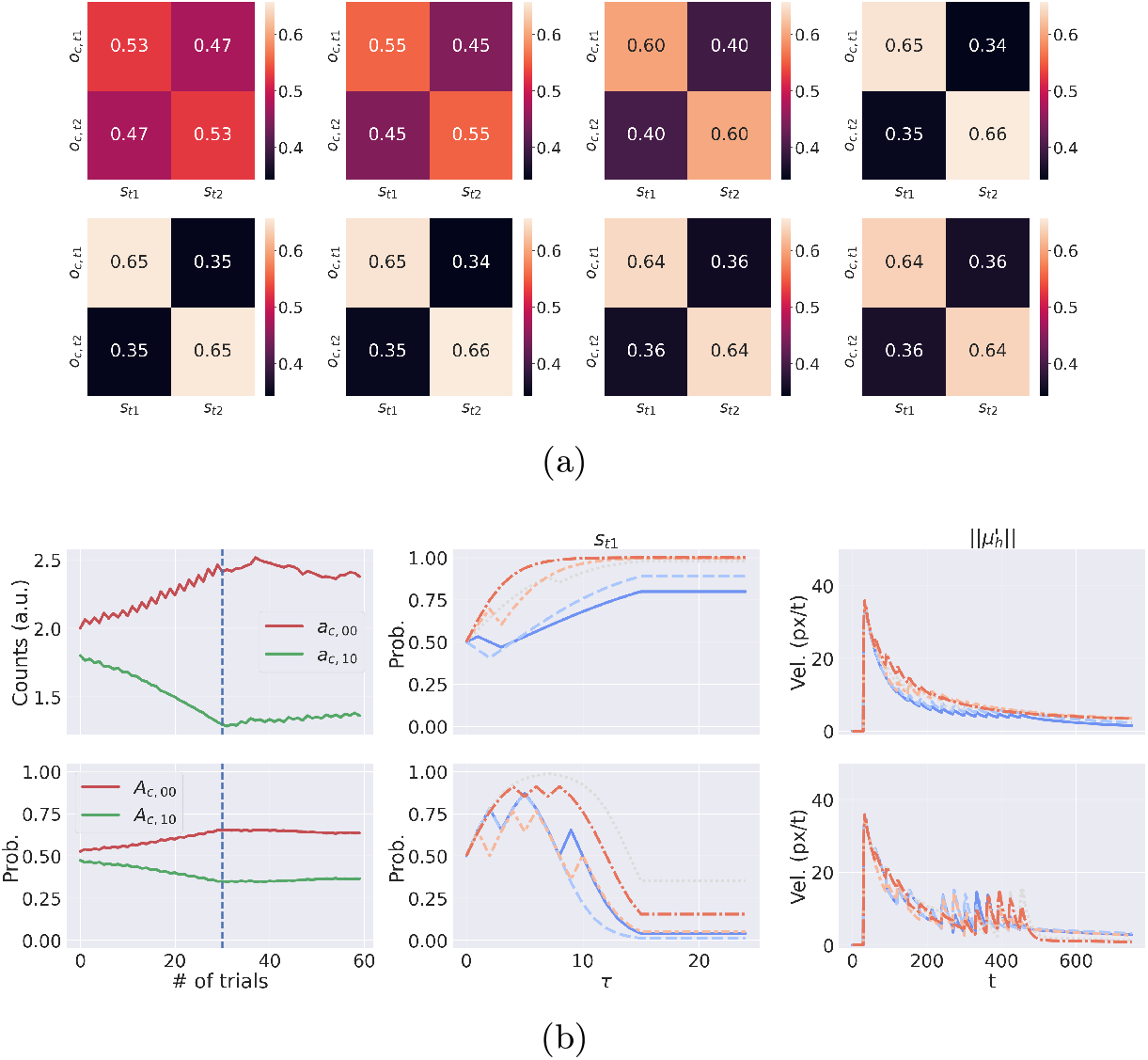
Results of Simulation 2. (a) Learning of ***A***_*c*_ in 30 congruent trials followed by 30 incongruent trials. The first (or second) row plots the values of ***A***_*c*_ for 4 equally spaced trials of the congruent (or incongruent) conditions. (b) Left column: state-outcomes coincidences ***a***_*c*_ (top), and likelihood matrix ***A***_*c*_ (bottom), for the left target. Middle column: discrete hidden states ***s*** of left target for 4 equally spaced congruent trials (top) and incongruent trials (bottom). Right column: norm of the estimated hand velocity 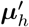 for 4 equally spaced congruent trials (top) and incongruent trials (bottom). Early-to-late trajectories are represented on a blue-to-red color scale. Note that for the sake of clarity we only plotted the dynamics associated with the left target, since the ones of the right target behave similarly.

### 3.3 Simulation 3: Learning a response strategy

Besides learning the mapping between targets and cues, the agent can also learn the mapping between targets and motor responses, affecting the adopted strategy (e.g., moving faster or slower, after observing a few or several cues). Changes in response strategies are common during cognitive experiments; for example, in the Posner task, participants often slow down their responses after encountering incongruent trials [19].

Recall that the discrete hidden states generate predictions ***o***_*h*_ related to three hand dynamics, i.e., reaching the left target, reaching the right target, or staying. In particular, the parameter *α*_*h*_ controls the strength of the third dynamics, and can be seen as the agent’s uncertainty over the strategy to adopt: the higher *α*_*h*_, the higher the probability of the correct target to initiate a movement. This is the consequence of the Bayesian model average over the three dynamics computed by Equation 8.

Learning the likelihood matrix ***A***_*h*_ involves counting the coincidences between target probabilities and hand dynamics, via Equation 12. This implies that if the agent’s hand is moving toward its target choice – as occurs more often during congruent trials – the related mapping will increase, leading to a decreased probability of the other dynamics, i.e. to stay or move to the right target. This is equivalent to a low uncertainty *α*_*h*_ which means that the agent’s urgency to move increases. Instead, if the agent spends significant time between the two targets – as occurs more often during incongruent trials – the mapping from the two targets to the third (stay) dynamics will increase. This means a high uncertainty *α*_*h*_, which translates to a low urgency to move.

To analyze this behavior, we ran a similar experiment to the previous one, i.e., 30 congruent trials followed by 30 incongruent trials. The counts ***a***_*h*_ are initialized to:

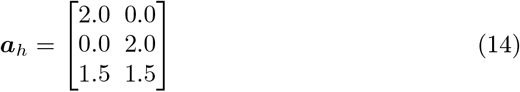

such that the parameter *α*_*h*_ is 0.43 – corresponding to a medium urgency to move. The forgetting and learning rates are respectively set to 0.99 and 0.01, while the parameter *α*_*c*_ is kept fixed to 0.4. As before, the correct choice is sampled from both targets for every trial. The results of this simulation are illustrated in Figure 4. During congruent trials, the third row of ***A***_*h*_ slightly decreases – meaning a lower strength of the stay dynamics; as shown in the top panel of the third column of Figure 4b – displaying the dynamics of the hand velocity – the increase in the agent’s confidence results in faster and stronger responses during late congruent trials (red trajectories). During incongruent trials, the learning is reversed and the last row of the matrix increases. In this case, the agent adapts its strategy to the new environmental uncertainty, and starts moving only when the correct cues are presented. Considering the bottom panel of the third column of Figure 4b, if early trials are characterized by two spikes of the estimated hand velocity 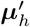 (corresponding to the initial movement toward the wrong target and the change of mind toward the correct one) at about *t* = 100 and *t* = 400, late trials only exhibit a single spike at about *t* = 500, with a much lower magnitude. This behavior is evident from Figure 4c, showing three sample trials during the incongruent phase; note how the agent learns a more cautious strategy, avoiding changes of mind in late trials. Finally, notice from the first column of Figure 4b that the learning of ***A***_*h*_ mainly involves modulation of the correct reaching movements and the stay dynamics, while the probability of opposite reaching movements (values on the antidiagonal) remains low.

**Fig. 4:**
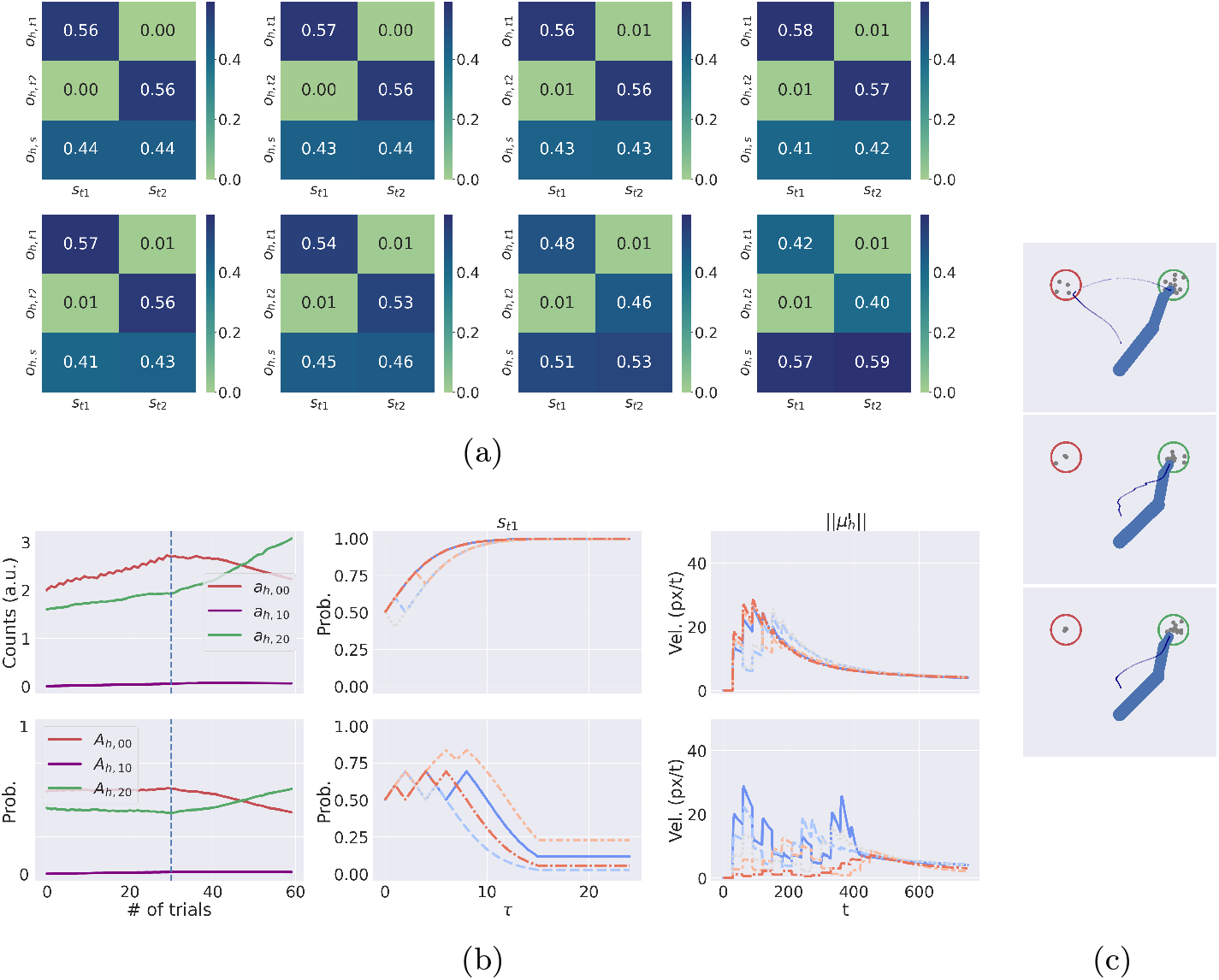
Results of Simulation 3. (a) Learning of ***A***_*h*_ in 30 congruent trials followed by 30 incongruent trials. The first (or second) row plots the values of ***A***_*h*_ for 4 equally spaced trials of the congruent (or incongruent) trials. (b) Left column: state-outcomes coincidences ***a***_*h*_ (top), and likelihood matrix ***A***_*c*_ (bottom), for the left target. Middle column: discrete hidden states ***s*** of left target for 4 equally spaced congruent trials (top) and incongruent trials (bottom). Right column: norm of the estimated hand velocity 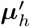 for 4 equally spaced congruent trials (top) and congruent trials (bottom). Early-to-late trajectories are represented on a blue-to-red color scale. (c) Agent’s trajectory (in dark blue) for early and late trials of incongruent conditions.

## 4 Discussion

Living organisms often face embodied decisions, which require not only selecting between outcomes but also simultaneously specifying, selecting between and executing plans to achieve these outcomes. Embodied decision models have begun to address these challenges by allowing decision and action processes to proceed in parallel and influence each other reciprocally [4,23]. A recent hybrid active inference model [35] successfully addresses embodied choices by using sensorimotor feedback from continuous dynamics – such as self-information [3] – as evidence for the decision itself.

In this study, we leverage and extend that model by studying how learning affects the agent’s behavior. In particular, we focused on three kinds of learning, namely, learning target priors, cue validity, and response strategies – all of which have been reported in human cognitive studies [21,29]. In the active inference model, these three types of learning correspond to keeping Dirichlet counts of the prior matrix ***D*** encoding target probabilities, of the likelihood matrix ***A***_*c*_ (i.e., the mapping between targets and cues), and of the likelihood matrix ***A***_*h*_ (i.e., the mapping between targets and hand dynamics).

Taken together, our simulations show that active inference agents can dynamically optimize the enaction of embodied decisions [28] based on the observed contexts. First, by learning the statistical structure of the task, the model can form priors about the correct target, which in turn determines fast and accurate movements even in case of conflicting sensory evidence. While previous studies showed that motor inference (i.e., inferring choice alternatives from action dynamics) helps optimize the speed-accuracy trade-off during embodied decisions [23,35], here we showed that it also affects the formation of habits, which in some cases can lead to incorrect task execution. Note however that incorrect choices generally lead to negative feedback, which might help overcome this problem during cognitive experiments. Second, we showed that the embodied decision model can successfully learn the likelihood mapping between observations (cues) and correct choice alternatives (targets). In turn, this allows the model to flexibly adapt to conditions in which cues are more or less informative, as in the case of experimental blocks with different cue validity in the Posner task [29]. Third, active inference models of embodied decision can flexibly adapt their response strategy and the urgency to move to task statistics – for example, by starting movement faster (or slower) in situations where decisions require fewer (or more) cues, as exemplified by congruent (or incongruent) trials.

Finally, our results illustrate the efficacy of learning discrete preferences and strategies using sensorimotor feedback from continuous dynamics. While a few studies addressed how to realize dynamic inference [31] and dynamic planning [33,34], learning in hybrid models of active inference is a yet unexplored topic. A promising research direction for the future would be to extend the present model with discrete transition distributions, and concurrently learn discrete and continuous dynamics, which might be key to advancing the use of active inference in tackling realistic tasks.

## Acknowledgments

This research received funding from the European Union’s Horizon 2020 Framework Programme for Research and Innovation under the Specific Grant Agreement No. 952215 (TAILOR); the European Research Council under the Grant Agreement No. 820213 (ThinkAhead), the Italian National Recovery and Resilience Plan (NRRP), M4C2, funded by the European Union – NextGenerationEU (Project IR0000011, CUP B51E22000150006, “EBRAINS-Italy”; Project PE0000013, CUP B53C22003630006, “FAIR”; Project PE0000006, CUP J33C22002970002 “MNESYS”), the PRIN PNRR P20224FESY, and the European Union’s Horizon H2020-EIC-FETPROACT-2019 Programme for Research and Innovation under Grant Agreement 951910. The GEFORCE Quadro RTX6000 and Titan GPU cards used for this research were donated by the NVIDIA Corporation. The funders had no role in study design, data collection and analysis, decision to publish, or preparation of the manuscript.

## References

1. Buckley, C.L., Toyoizumi, T.: A theory of how active behavior stabilises neural activity: Neural gain modulation by closed-loop environmental feedback. PLoS computational biology 14(1), e1005926 (2018)

2. Burk, D., Ingram, J.N., Franklin, D.W., Shadlen, M.N., Wolpert, D.M.: Motor effort alters changes of mind in sensorimotor decision making. PLoS ONE 9(3), e92681 (Mar 2014). 10.1371/journal.pone.0092681

3. Chen, C.L., Aymanns, F., Minegishi, R., Matsuda, V.D., Talabot, N., Günel, S., Dickson, B.J., Ramdya, P.: Ascending neurons convey behavioral state to integrative sensory and action selection brain regions. Nature neuroscience 26(4), 682–695 (2023)

4. Christopoulos, V., Schrater, P.R.: Dynamic integration of value information into a common probability currency as a theory for flexible decision making. PLoS computational biology 11(9), e1004402 (2015)

5. Cisek, P.: Cortical mechanisms of action selection: the affordance competition hypothesis. Philosophical Transactions of the Royal Society B: Biological Sciences 362(1485), 1585–1599 (2007)

6. Cisek, P.: Making decisions through a distributed consensus. Current opinion in neurobiology 22(6), 927–936 (2012)

7. Cisek, P., Kalaska, J.F.: Neural mechanisms for interacting with a world full of action choices. Annual Review of Neuroscience 33, 269–298 (2010). 10.1146/annurev.neuro.051508.135409

8. Cisek, P., Pastor-Bernier, A.: On the challenges and mechanisms of embodied decisions. Philosophical Transactions of the Royal Society B: Biological Sciences 369(1655), 20130479 (2014)

9. Cisek, P., Puskas, G.A., El-Murr, S.: Decisions in changing conditions: The urgency-gating model. Journal of Neuroscience 29(37), 11560–11571 (2009). 10.1523/JNEUROSCI.1844-09.2009

10. Cos, I., Pezzulo, G., Cisek, P.: Changes of mind after movement onset depend on the state of the motor system. Eneuro 8(6) (2021)

11. Da Costa, L., Parr, T., Sajid, N., Veselic, S., Neacsu, V., Friston, K.: Active inference on discrete state-spaces: A synthesis. Journal of Mathematical Psychology 99 (2020). 10.1016/j.jmp.2020.102447

12. Eriksen, B.A., Eriksen, C.W.: Effects of noise letters upon the identification of a target letter in a nonsearch task. Perception and Psychophysics 16(1), 143–149 (Jan 1974). 10.3758/bf03203267

13. Eriksen, C.W., Schultz, D.W.: Information processing in visual search: A continuous flow conception and experimental results. Perception & psychophysics 25(4), 249–263 (1979)

14. Freeman, J.B., Dale, R., Farmer, T.A.: Hand in motion reveals mind in motion. Frontiers in psychology 2, 59 (2011)

15. Friston, K., FitzGerald, T., Rigoli, F., Schwartenbeck, P., O’Doherty, J., Pezzulo, G.: Active inference and learning. Neuroscience and Biobehavioral Reviews 68, 862–879 (Sep 2016). 10.1016/j.neubiorev.2016.06.022

16. Friston, K., Parr, T., Zeidman, P.: Bayesian model reduction pp. 1–32 (2018), http://arxiv.org/abs/1805.07092

17. Friston, K., Penny, W.: Post hoc Bayesian model selection. NeuroImage 56(4), 2089–2099 (2011). 10.1016/j.neuroimage.2011.03.062

18. Friston, K.J., Parr, T., de Vries, B.: The graphical brain: Belief propagation and active inference 1(4), 381–414 (2017). 10.1162/NETN, 10.1162/NETN_a_00018

19. Gómez, C.M., Arjona, A., Donnarumma, F., Maisto, D., Rodríguez-Martínez, E.I., Pezzulo, G.: Tracking the time course of bayesian inference with event-related potentials: A study using the central cue posner paradigm. Frontiers in Psychology 10, 1424 (2019)

20. Gordon, J., Maselli, A., Lancia, G.L., Thiery, T., Cisek, P., Pezzulo, G.: The road towards understanding embodied decisions. Neuroscience & Biobehavioral Reviews 131, 722–736 (2021)

21. Gratton, G., Coles, M.G., Donchin, E.: Optimizing the use of information: strategic control of activation of responses. Journal of Experimental Psychology: General 121(4), 480 (1992)

22. Grießbach, E., Raßbach, P., Herbort, O., Cañal-Bruland, R.: Embodied decisions during walking. Journal of Neurophysiology 128(5), 1207–1223 (2022)

23. Lepora, N.F., Pezzulo, G.: Embodied choice: How action influences perceptual decision making. PLOS Computational Biology 11(4), e1004110 (Apr 2015). 10.1371/journal.pcbi.1004110

24. Marcos, E., Cos, I., Girard, B., Verschure, P.F.: Motor cost influences perceptual decisions. PLoS One 10(12), e0144841 (2015)

25. Parr, T., Friston, K.J.: The Discrete and Continuous Brain: From Decisions to Movement—And Back Again Thomas. Neural Computation 30, 2319–2347 (2018). 10.1162/neco_a_01102

26. Parr, T., Pezzulo, G., Friston, K.J.: Active inference: the free energy principle in mind, brain, and behavior (2022)

27. Pezzulo, G., Cisek, P.: Navigating the Affordance Landscape: Feedback Control as a Process Model of Behavior and Cognition. Trends in Cognitive Sciences 20(6), 414–424 (2016). 10.1016/j.tics.2016.03.013

28. Pezzulo, G., Donnarumma, F., Iodice, P., Maisto, D., Stoianov, I.: Model-based approaches to active perception and control. Entropy 19(6) (2017). 10.3390/e19060266

29. Posner, M.I.: Orienting of attention. Quarterly journal of experimental psychology 32(1), 3–25 (1980)

30. Priorelli, M., Pezzulo, G., Stoianov, I.: Active vision in binocular depth estimation: A top-down perspective. Biomimetics 8(5) (2023). 10.3390/biomimetics8050445

31. Priorelli, M., Stoianov, I.: Dynamic inference by model reduction. bioRxiv (2023). 10.1101/2023.09.10.557043

32. Priorelli, M., Pezzulo, G., Stoianov, I.P.: Deep kinematic inference affords efficient and scalable control of bodily movements. Proceedings of the National Academy of Sciences of the United States of America 120 (2023). 10.1073/pnas.2309058120

33. Priorelli, M., Stoianov, I.P.: Deep hybrid models: infer and plan in the real world. arXiv (2024). 10.48550/arXiv.2402.10088

34. Priorelli, M., Stoianov, I.P.: Dynamic planning in hierarchical active inference. arXiv (2024). 10.48550/arXiv.2402.11658

35. Priorelli, M., Stoianov, I.P., Pezzulo, G.: Embodied decisions as active inference. bioRxiv (Jun 2024). 10.1101/2024.05.28.596181

36. Ratcliff, R., McKoon, G.: The diffusion decision model: Theory and data for two-choice decision tasks. Neural Computation 20(4), 873–922 (Apr 2008). 10.1162/neco.2008.12-06-420

37. Smith, R., Friston, K.J., Whyte, C.J.: A step-by-step tutorial on active inference and its application to empirical data. Journal of Mathematical Psychology 107, 102632 (2022). 10.1016/j.jmp.2021.102632

38. Song, J.H., Nakayama, K.: Hidden cognitive states revealed in choice reaching tasks. Trends in cognitive sciences 13(8), 360–366 (2009)

39. Spivey, M.: The continuity of mind. Oxford University Press (2008)

40. Stoianov, I., Maisto, D., Pezzulo, G.: The hippocampal formation as a hierarchical generative model supporting generative replay and continual learning. Progress in Neurobiology 217, 1–20 (2022). 10.1016/j.pneurobio.2022.102329

41. Usher, M., McClelland, J.L.: The time course of perceptual choice: the leaky, competing accumulator model. Psychological review 108(3), 550 (2001)

42. Wispinski, N.J., Gallivan, J.P., Chapman, C.S.: Models, movements, and minds: bridging the gap between decision making and action. Annals of the New York Academy of Sciences 1464(1), 30–51 (2020)

43. Ye, W., Damian, M.F.: Effects of conflict in cognitive control: Evidence from mouse tracking. Quarterly Journal of Experimental Psychology 76(1), 54–69 (2023)

44. Yoo, S.B.M., Hayden, B.Y., Pearson, J.M.: Continuous decisions. Philosophical Transactions of the Royal Society B 376(1819), 20190664 (2021)

